# Maturation of Neuronal Activity in Caudalized Human Brain Organoids

**DOI:** 10.1101/779355

**Authors:** Svetlana M. Molchanova, Mariia Cherepkova, Shamsiiat Abdurakhmanova, Suvi Kuivanen, Elina Pörsti, Päivi A. Pöhö, Jaakko S. Teppo, Jonna Saarimäki-Vire, Jouni Kvist, Diego Balboa, Andrii Domanskyi, Raul R. Gainetdinov, Petteri Piepponen, Markku Varjosalo, Risto Kostiainen, Olli Vapalahti, Tomi P. Taira, Timo Otonkoski, Maxim M. Bespalov

**Affiliations:** Stem Cells and Metabolism Research Program, Faculty of Medicine, University of Helsinki, Finland; Neuroscience Center, HiLIFE Helsinki Institute of Life Science, University of Helsinki, Finland; Department of Anatomy, University of Helsinki, Helsinki, Finland; Department of Virology, Medicum, University of Helsinki, Helsinki, Finland; Drug Research Program and Division of Pharmaceutical Chemistry and Technology, Faculty of Pharmacy, University of Helsinki, Helsinki, Finland; Institute of Biotechnology, HiLIFE Helsinki Institute of Life Science, University of Helsinki, Finland; Institute of Translational Biomedicine and Saint Petersburg University Hospital, Saint Petersburg State University, Saint Petersburg, Russia; Department of Veterinary Biosciences, University of Helsinki, Helsinki, Finland; Division of Clinical Microbiology, University of Helsinki and Helsinki University Hospital, Helsinki, Finland; Children’s Hospital, Helsinki University Hospital and University of Helsinki, Helsinki, Finland; Faculty of Bio- and Environmental Sciences, Molecular and Integrative Biosciences Research Program; Department of Biosystems Science and Engineering, ETH Zurich, Basel, Switzerland; Novo Nordisk Research Center Oxford, Oxford, United Kingdom; Centre for Genomic Regulation, The Barcelona Institute of Science and Technology, Barcelona, Spain

## Abstract

Human brain organoids are an emerging tool to study functional neuronal networks in health and disease. A critical challenge is the engineering of brain organoids with defined regional identity and developmental stage. Here we describe a protocol for generating hindbrain-like organoids from human pluripotent stem cells. We first generated a stable pool of caudalized stem cells that expressed hindbrain identity transcription factors and differentiated into tissue containing neurons and astrocytes. After maturation, caudalized brain organoids presented synaptically connected networks consisting of glutamate-, GABA-, and serotoninergic postmitotic neurons. These mature neurons displayed electric properties and dendritic trees resembling medulla oblongata neurons. They fired spontaneous and evoked repetitive action potentials, released serotonin and displayed excitatory and inhibitory synaptic currents, functionally resembling the activity patterns observed in normal human fetal brain. Reminiscent of infected human fetal brain, infection with Zika virus hampered organoid development, while the treatment with anticonvulsant drugs - carbamazepine and valproic acid - reduced organoid growth. Neuronal maturation also occurred in the grafted organoids *in vivo*. In conclusion, our approach enables efficient derivation of caudalized neuronal stem cells that differentiate into mature and functional neurons in organoids with hindbrain identity following human developmental trajectory. The organoids provide excellent model to study congenital abnormalities in brain development and for drug testing.

## Introduction

Understanding human brain functioning and development, as well as utility of this knowledge in drug discovery has been hampered by the availability of living human brain tissue. Many aspects of the normal and diseased human brain can be modelled in animals, but the species-specific differences can lead to underwhelming modeling results. One of the prominent structural differences between mouse and human brain, besides a sheer size, is the gyration that is believed to result from the increased expansion potential of the radial glia cells residing in a differently structured subventricular zone (Lui et al., 2011). Increased cell diversity is also a factor to consider when modeling human brain in other species. For example, some inhibitory neurons are primate- or human-specific (Yáñez et al., 2005; Luo et a., 2017; Boldog et al., 2018). In drug discovery, marked differences in receptor affinities and cellular responses to small molecules in model animals can negate their utility as drugs in human patients (Hu et al., 2009). Better understanding of the differences between human and animal brain and development of *in vitro* platforms for drug testing justifies attempts to create more efficient techniques for human nervous tissue generation from stem cells.

Over the past decade, different methods have been developed to produce brain-like tissue, organoids, from stem cells. Cells in brain organoids self-organize into structured and functional nervous tissue, providing insights into the development, morphology and evolution of human brain, and for disease modelling (Eiraku et al., 2008; Lancaster et al., 2013; Mariani et al., 2015; Mora-Bermúdez et al., 2016). Pioneering studies have proved that neuronal migration, supported by radial glia and formation of cortical layers can be modelled *in vitro* using 3D systems (Lancaster et al., 2017, Birey et al., 2017; Xiang et al., 2017). Brain organoids provide a unique possibility to study early steps of human neuronal network development and to compare them with that in animal models. Establishment of synaptic connectivity and the development of cortical neurons oscillatory activity have already been described both in 2D (Kirwan et al., 2015, Kuijlaars et al., 2016, Mäkinen et al., 2018) and 3D models (Li et al., 2017, Trujillo et al., 2019).

However, neurons directly derived from human pluripotent stem cells (hPSC) often fail to reach maturity and functionality, and producing specific neuronal subtypes is often challenging (Wu & Hochedlinger, 2011). Brain organoids that develop directly from hPSC contain cells with different regional identity, including sensory epithelium cells. Directly differentiated whole-brain organoids are thus very heterogeneous with unpredictable morphology and internal structure. They also require a long differentiation time (e.g. astrocytes appear after 3 months; Quadrato et al., 2017). One approach to solve this problem is to pre-pattern stem cells and use forced differentiation by modulating specific regionalization pathways. Generation of brain organoids from expandable and stable pool of neural stem cells offers better homogeneity of starting cell population, faster differentiation, and a lack of non-neural components (Monzel et al., 2017). Various regionalized organoids were generated from pre-patterned stem cells: dorsal and ventral forebrain-, hippocampus-, cerebellum- and midbrain-like organoids (Paşca, 2018).

To date, there were no attempts made yet to model hindbrain structures. Hindbrain contains pons, cerebellum (both derived from metencephalon), and medulla (from a more caudal myelencephalon). Together they control critical autonomic functions such as respiration, heartbeat as well as motor activity, sleep and wakefulness. Here, we aimed to model hindbrain development from pre-patterned primitive neural stem cells. To this end, we optimized a protocol for generation of caudalized human neuroepithelial stem cells (hNESC) from both ES and induced PSC, and generated hindbrain-like organoids, containing functional neurons organized into developing networks. We performed metabolomic and proteomic profiling of these cells as well as targeted gene expression analysis. We studied the development of neuronal activity pattern in organoids of different ages, recapitulating some of hindbrain’s early developmental hallmarks and probed the applicability of our model for studying infectious diseases and for detecting drugs teratogenicity.

## Results

### Human neuroepithelial stem cells generation

Neuroepithelial cells are the proliferating stem cells of developing brain that were formed from pseudostratified epithelium. NESC give rise to radial glia progenitor cells that in turn differentiate into neurons and glia. We tested whether we can establish a stable population of caudalized NESC and differentiate them into neurons.

Human PSC were differentiated to NESC via embryoid bodies with neural lineage specification induced by inhibition of transforming growth factor β (TGFβ) and bone morphogenic protein (BMP), and by induction of WNT and Hedgehog signaling pathways (Fig. 1A). Compared to published protocols of neural induction (Reinhardt et al., 2013) we observed less cell death and higher yield of hNESC (Supplementary fig. 1A).

**Figure 1.**
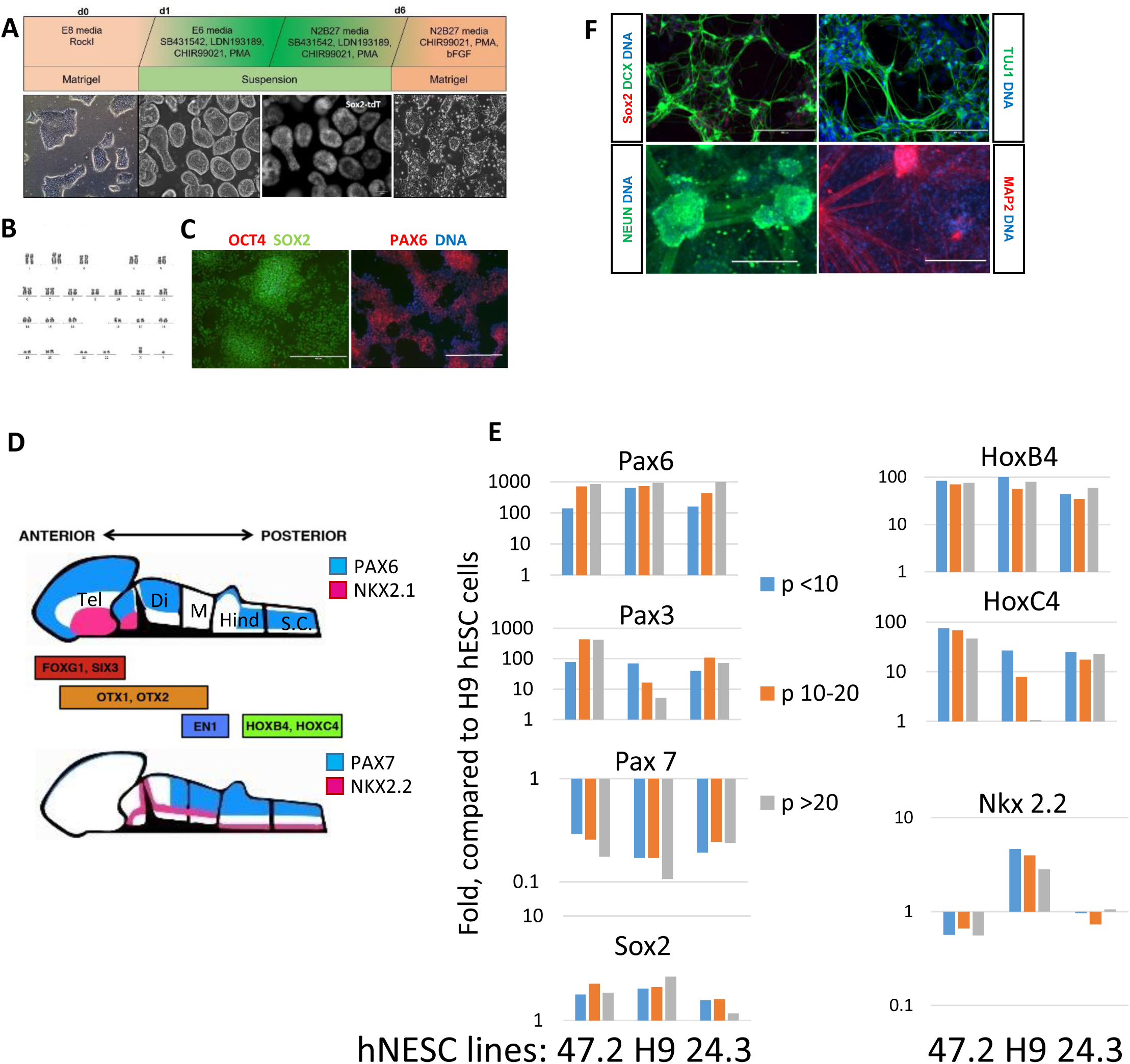
Human neuroepithelial stem cell (hNESC) generation and characterization. **A**. hNESC generation protocol scheme. RockI – ROCK inhibitor; PMA – purmorphamine; Sox2-tdT – SOX2-tdTomato reporter direct fluorescence (Balboa et al., 2017). **B**. Passage 20 (P20) hNESC are of a normal karyotype. **C**. Master pluripotency gene OCT4 is downregulated in hNESC, while SOX2 and PAX6 are expressed. Bar is 400 µm. **D**. Expression of neuronal markers in developing neural tube *in vivo* (modified from Imaizumi et al., 2015). **E**. Quantitative RT-PCR of neuronal genes in three hNESC lines derived from HEL47.2 (47.2); H9 hESC (H9); HEL24.3 (24.3) at various passages (p) expressed as fold change of expression in parental (pluripotent) H9 hESC line. **F**. Neuronal differentiation on hNESC in two-dimensions. Bar is 200 µm in upper panels and 400 µm in a lower panel. **E**.

After plating, the passage (p) zero colonies already lacked OCT4 and GATA4 but expressed SOX2, VIMENTIN, occasionally ASCL1 (MASH1) and doublecortin (DCX) – one of the early neural markers (Supplemental fig. 1B). During passaging hNESC formed compact colonies of rapidly proliferating cells that were cultured for up to 45 passages without notable changes in morphology, gene expression profile and maintaining normal karyotype (p20; Fig. 1B, E). Immunostaining of later passage hNESC demonstrated nearly uniform distribution of SOX2 and PAX6 transcription factors and the absence of master pluripotency gene OCT4 (Fig. 1C).

To determine identity of newly formed NESC, we analyzed the expression of selected region-specific genes (Fig. 1D). The qPCR data demonstrated a downregulation of *FOXG1* (*NKX2.1* did not change significantly; not shown), and upregulation of medial *PAX6* but not of the dorsal *PAX7* marker and induction of some ventro-caudal markers – *MSX1* (floor plate), *HOXA2*, *HOXB4* and caudal *HOXC4* (Fig. 1E; Supplementary fig. 1C). However, no upregulation of *OTX2* (caudal forebrain and periventricular cells of the mesencephalon) was observed (not shown). (Fig. 1E; Supplementary fig. 1C) (Imaizumi et al., 2015; Fig. 1D). Another ventral marker *NKX2.2* was moderately (3.8 folds) upregulated but only in one hNESC line (H9-ESC-derived). We also observed a variable level of *PAX3* expression in different passages, which could be explained by the presence of a subpopulation of neural crest stem cells (Reinhardt et al., 2013). Based on these data, we concluded that most hNESC had ventromedial hindbrain identity (Imaizumi et al., 2015).

Upon withdrawal of mitogens, hNESCs underwent spontaneous differentiation into doublecortin-positive neural progenitors and then into more mature neurons (Fig. 1F, 5G). This could be facilitated by NOTCH-signaling inhibitor, cAMP analog, and brain-derived neurotrophic factor (BDNF) – forced differentiation, for future reference.

Proteomic profiling detected marked downregulation of pluripotency related proteins (DNMT3A/B, DPPA4, and L1TD1) in NESC compared to hPSC. Conversely, a radial glial marker FABP7 and COX6A1 were upregulated the most. In neural progenitors (forced differentiation of hNESC for 3 days, Supplemental fig. 1D) doublecortin, internexin and nestin (markers of neuroblasts and of young neurons) were upregulated as well as neuromodulin (GAP43) and MAP2 (markers of more mature neurons) (Supplementary table 1). We also performed untargeted metabolic profiling in the same cells and the principal component analysis (PCA) has clearly separated all three groups (Supplemental Fig. 1E).

Although we successfully generated hNESC with presumed caudalized ventromedial identity the prolonged differentiation in 2D resulted in aggregate formation (Fig. 1F) and spontaneous detachment. This precluded further maturation of hNESC-derived neurons in 2D and prompted us to develop a 3D organoid culture system.

### Brain organoids generation and identity

To improve maturation of neurons from caudalized NESC we aggregated them (on average 5 × 10^3^ cells) in round-bottom ultra-low-adhesion 96-well plates. For the first 6 days, we grew spheroids in a standard culture medium, containing WNT- and Hedgehog-pathways stimulants along with basic FGF. On day 7 aggregates were transferred to the orbital shaker into the media devoid of stimulants and FGF. A test of optimal culture conditions revealed that WNT-signaling pathway stimulation increased organoid growth the most (Fig. 2B).

**Figure 2.**
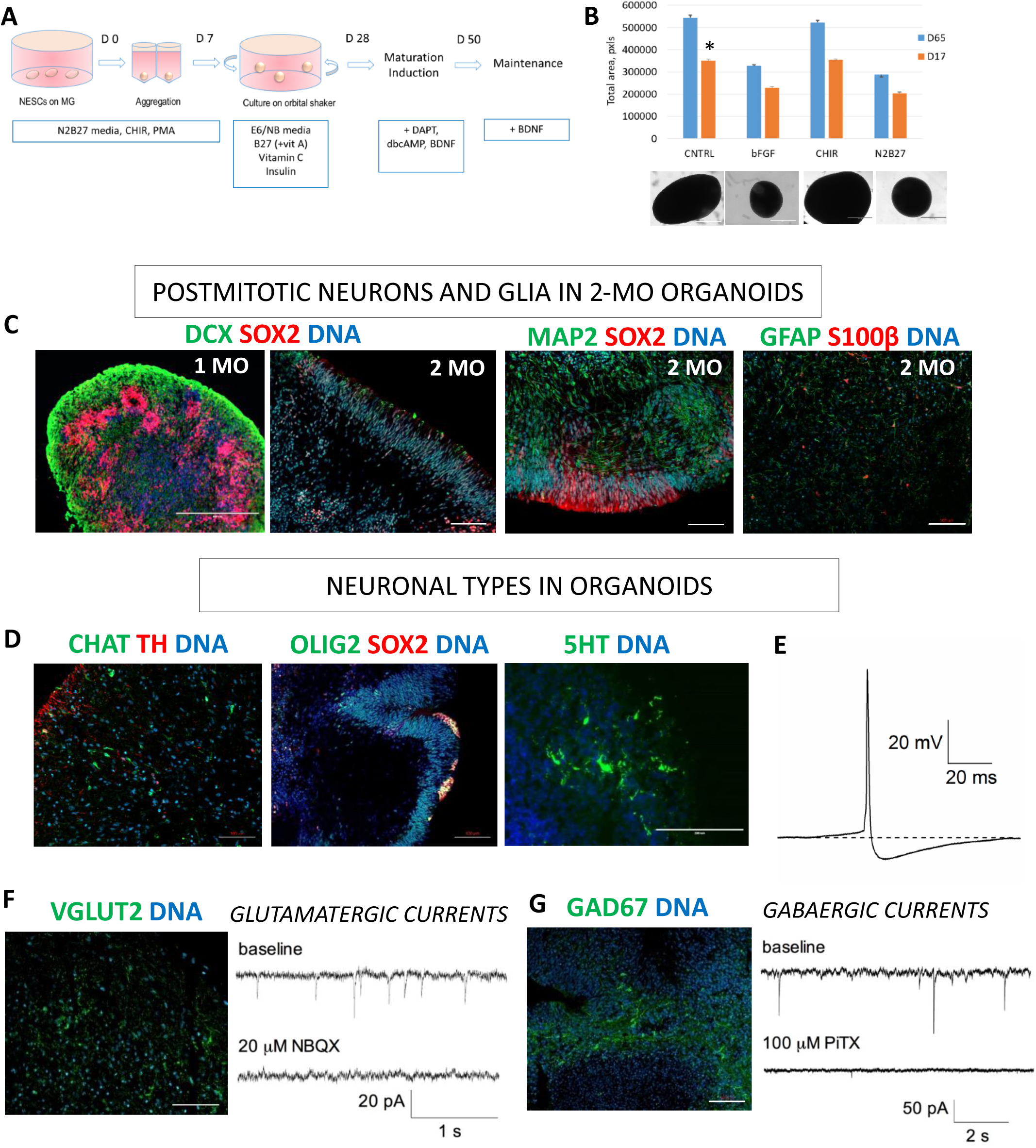
Brain organoids generation, growth and expression of neuronal markers. **A**. A scheme for generating brain organoids from hNESC. **B**. Role of FGF and Wnt signaling in organoid growth. Control, full media (Cntrl); basic FGF (FGF2, bFGF); CHIR99021 (CHIR); base media - N2B27; days post aggregation (D). *p<0,05, two-way ANOVA. **C**. Neural stem cells marker (SOX2) and a marker of young neurons doublecortin (DCX) are downregulated in a process of maturation induction. In 2-month-old organoids there is a prominent expression of postmitotic neuron marker MAP2 and of astrocytic markers GFAP and S100β. **D**. Some neuronal precursors in 2-month-old organoids express motoneuron marker OLIG2, along with SOX2, and organoids contain choline acetyltransferase (ChAT)-positive and serotonergic (5HT) neurons but low level of TH-positive neurons. **E**. Action potential waveform of a neuron from 4-month-old organoid. Large action potential and long after hyperpolarization are characteristic for serotonergic neurons. **F**. Several neurons express vesicular glutamate transporter (VGLUT2) and exhibit spontaneous synaptic sodium currents, inhibited with AMPA receptor antagonist NBQX. **G**. Some neurons express glutamate decarboxylase (67 kDa isoform, GAD67), a marker of GABAergic neurons, and have spontaneous synaptic chloride currents that can be inhibited with GABAA blocker picrotoxin (PiTX).

During the first month of differentiation, organoids increased in size and presented abundant SOX2- and DCX-positive neuronal progenitors (Fig. 2C). To increase differentiation efficiency, on day 28 we changed culture medium to forced differentiation media in which organoids were cultured for 4 weeks (Fig. 2A).

Further incubation of brain organoids (2 to 4 months) resulted in neuronal progenitor markers (DCX and SOX2) decrease and MAP2 and GFAP immunoreactivity emergence (Fig. 2C; Supplemental movie 1), indicating the presence of both neurons and astroglia. Remaining progenitors tend to localize in periphery of the organoids (Fig. 2C, D). Compared to published protocols (Lancaster et al., 2013; Quadrato et al., 2017) we detected astrocytic maturation, characterized by expression of not only GFAP but also S100β, already in 2-month-old organoids, suggesting a higher efficiency of glia differentiation in organoids derived directly from pre-patterned hNESC (Fig. 2C).

Caudalized hNESC express HOXA2 and HOXB4 and are thus likely to differentiate into hindbrain’s *raphe nuclei* cells. Consistent with this, organoids produced serotonin (0.6-2.9 ng/ml; Supplemental fig. 2) and the presence of serotonergic neurons was confirmed by immunostaining (Fig. 2D). Gene expression, serotonin production, along with a relatively small number of dopaminergic cells identified by tyrosine hydroxylase (TH) labeling (Fig. 2D) suggests an identity closer to *rostral ventromedial medulla* (RVM), rather than to more rostral *dorsal raphe nucleus* (Ikemoto, 2007; Lu et al., 2016). In line with these findings, in mature organoids we also found choline-acetyltransferase- and OLIG2-positive cells, which may correspond to medullar motor neurons and their precursors (Fig. 2D)(Spinella et al, 1997).

To further investigate the hindbrain identity, we analyzed passive membrane properties and evoked action potentials in 4-month-old organoids. Four-month-old neurons had relatively small membrane capacitance and high input resistance (Table 1, Fig. 3B, C). Some neurons fired spontaneous action potentials with high frequency, characteristic for serotonergic neurons of *pons* and *medulla oblongata* (Table 1; Scott et al., 2005; Lu et al., 2016). Upon injection of a depolarizing current, most cells fired large-amplitude action potentials with long after-hyperpolarization, which is also similar to RVM, which includes serotonergic *nucleus raphe magnus* (Fig. 2E, Table 1). In addition, neurons in 4-month-old organoids had aspiny dendrites with small arborization, similar to neurons of RVM (Fig. 3D; Winkler et al., 2006).

**Figure 3.**
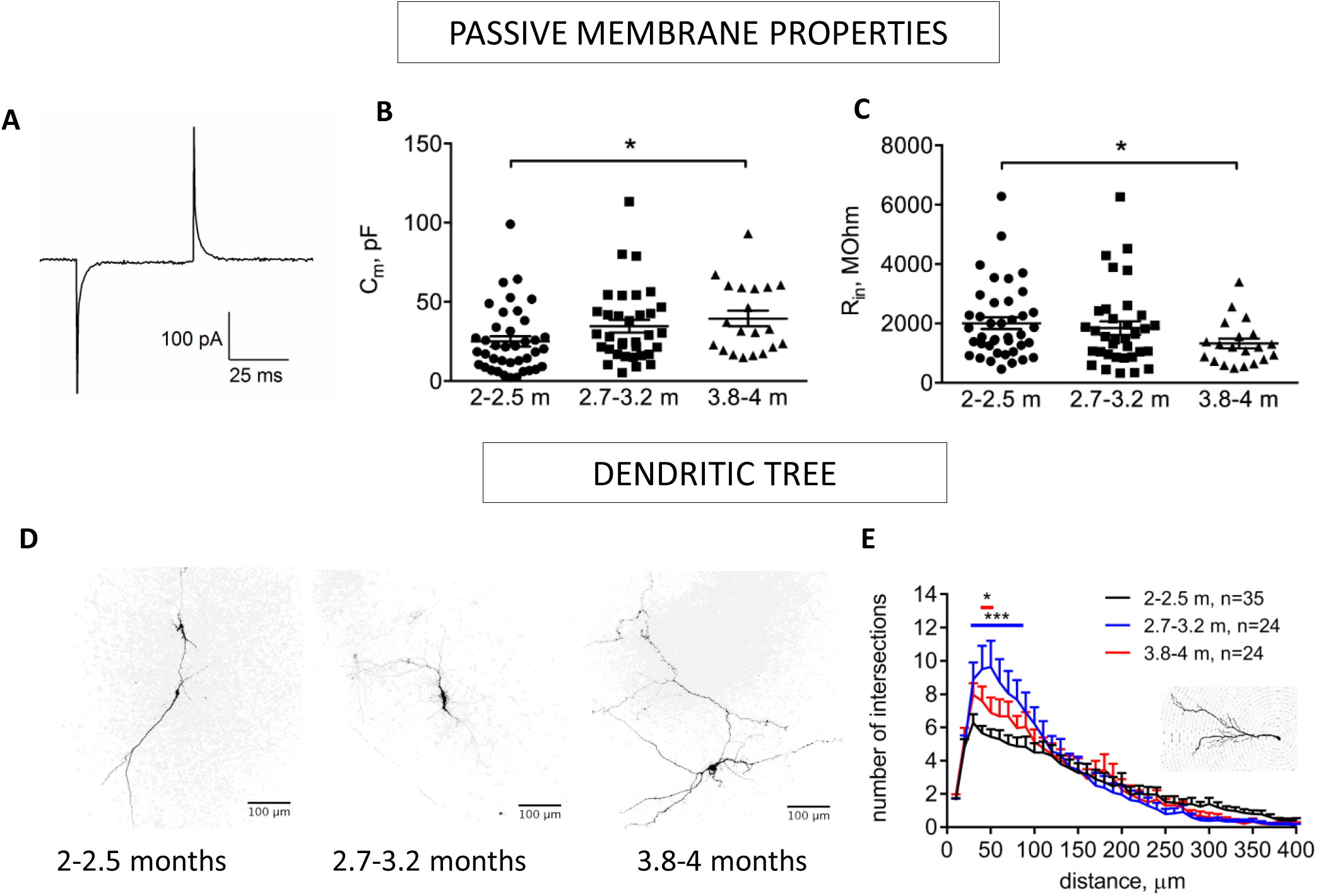
Passive membrane properties of the patched neurons (A-C) and their dendritic tree development over time (D-E). **A**. Traces showing current response to 5 mV voltage step in a voltage-clamped (-70 mV) cell. **B**. Membrane capacitance of voltage-clamped cells in organoids of various ages. Months – (m). **C**. Input resistance of voltage-clamped cells in organoids of various ages. *p<0,05, one-way ANOVA. **D**. Dendritic tree development of biocytin-filled cells in organoids of various ages. **E.** Sholl analysis of dendritic tree in neurons of different ages. Results are expressed as mean+SEM. *p<0,05 (2-month-old vs 4-month-old cells); ***p<0,001 (2-month-old vs 3-month-old cells), two-way ANOVA with Tukey’s multiple comparisons test.

**Table 1.** Passive membrane and firing properties in 4-month-old organoids resemble characteristics of serotonergic neurons

|  | Our data | Serotonergic neurons, differentiated from hiPSCs (Lu et al., 2016) |
| --- | --- | --- |
| Capacitance | 46±5.6 pF (n=20) | 25.0±1.8 pF |
| Spontaneous firing frequency | 2.0±0.7 Hz (35 %, 7 of 20 cells recorded) | 2.9±0.2 Hz |
| Action potential threshold | -35.9±1.0 mV (n=15) | -30 mV |
| Action potential amplitude | 64.9±2.2 mV (n=15) | 65.3±1.0 mV |
| Action potential half-width | 1.5±0.1 mV (n=15) |  |
| After hyperpolarization amplitude | 6.8±0.9 mV (n=15) | 14.3±0.6 mV |
| After hyperpolarization half-width | 40.6±6.4 ms (n=15) |  |

Organoids also contained glutamatergic and GABAergic neurons, shown by the immunostaining against VGLUT2 and GAD67 (Fig. 2F, G). The presence of these cell types was confirmed by recording AMPA receptor- and GABAA receptor-mediated synaptic currents. Glutamate currents were inhibited with AMPA receptor inhibitor (20 μM NBQX), while chloride currents were almost completely inhibited by 100 μM picrotoxin, a GABAA receptor inhibitor. The remaining currents were mediated perhaps by glycine receptors, as glycinergic synaptic transmission has also been shown in ventromedial medulla (Hossaini et al., 2012) (Fig. 2G).

Altogether, our immunostaining and functional data show that brain organoids contain serotonergic, cholinergic, GABAergic and glutamatergic neurons, low level of dopaminergic neurons, the neuronal passive and active membrane properties along with cellular morphology correspond to hindbrain that is in accordance with the gene expression data in hNESCs.

### Functional maturation

During neuronal growth and maturation the membrane capacitance increases, primarily due to dendritic growth, and conductivity decreases, due to increased insertion of ion channels into plasma membrane. Indeed, in 2- to 4-month-old brain organoids the membrane capacitance increased steadily, indicating the cell size growth (dendritic tree and its complexity, Fig. 3B, D, E). Input resistance, conversely, decreased with age of the organoids (Fig. 3C). In addition, the resting membrane potential decreased with age and was close to -50 mV in 4-month-old neurons, suggesting functional neuronal maturation (Fig. 4D). This development of passive membrane properties occurred together with the growth of dendritic tree (Fig. 3D, E). The Sholl analysis (a method to quantify the number of branches at variable distances from soma) revealed that the highest number of intersections with concentric circles was in soma’s proximity and in 3-month-old organoids. This number decreased in 4-month-old organoids, reflecting the disappearance of temporary short dendrites.

**Figure 4.**
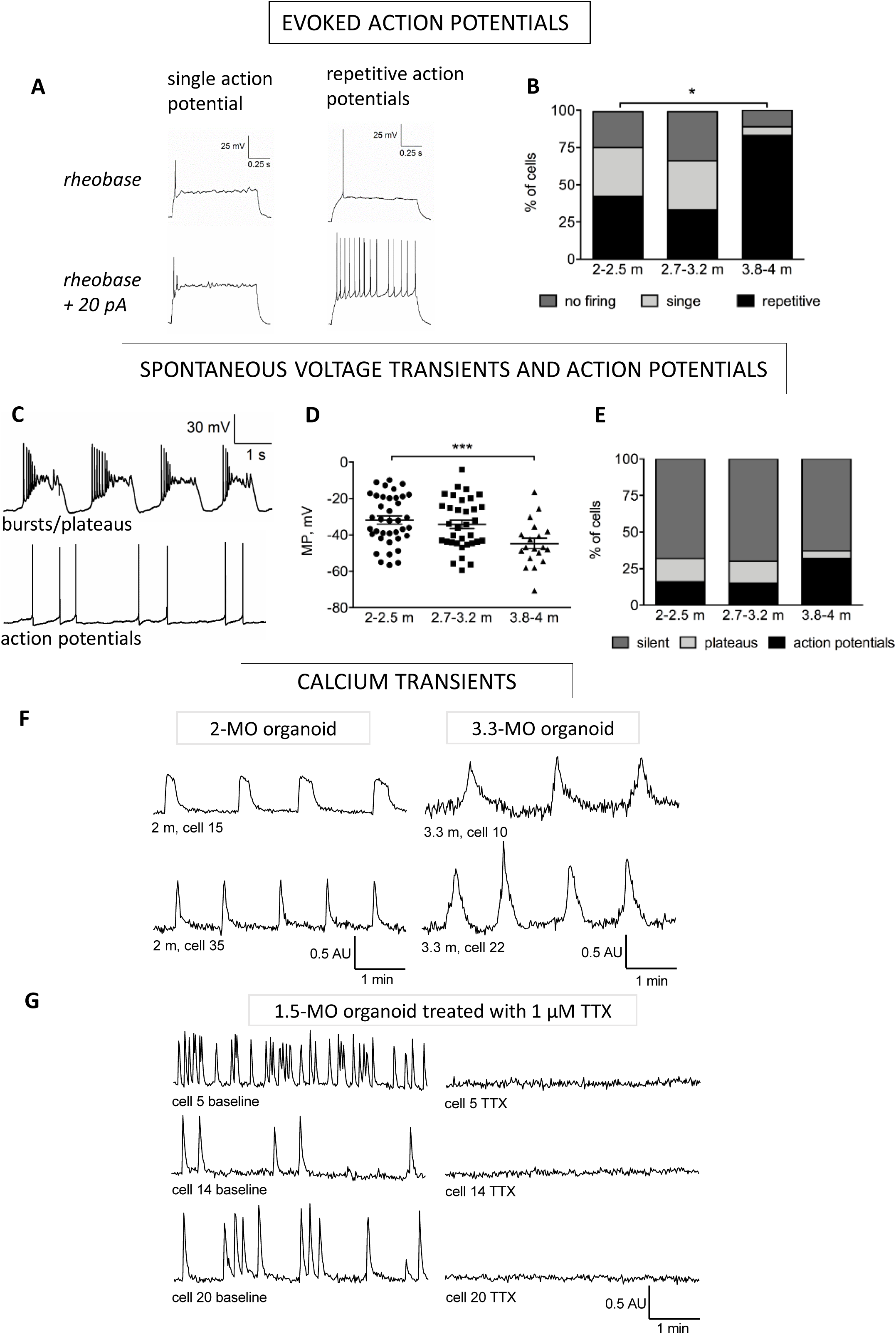
Development of the action potential firing and spontaneous electrical activity, and calcium oscillations in neurons of different ages. **A**. Example traces of evoked action potentials recorded after injecting depolarizing current steps (10 pA increment) for one second. **B**. Percentage of cells exhibiting repetitive evoked action potentials increases significantly in 4-month-old organoids. *p<0,05, Chi-square test. **C**. Example traces of spontaneous plateau-like depolarizations and action potentials. **D**. Membrane potential of cells in organoids of different ages, ***p<0,001 by one-way ANOVA. Months – (m). **E**. Percentage of cells in organoids of different ages, exhibiting different activity patterns. **F**. Variable forms of calcium transients recorded with Fluo4 dye in 2 and 3 months old (MO) organoids. **G**. Example traces from an organoid 17 days after forced induction of differentiation before (left) and after (right) application of 1 µM tetrodotoxin (TTX).

Ability to generate action potentials and to form synaptic connections are the main features of functional postmitotic neurons. The first brain organoid neurons that fired action potential started to appear in two-month-old organoids (Fig. 4A, B). However, only 42% (14 out of 33 cells) were able to generate repetitive action potentials in response to depolarizing current injection. Other cells either generated only single action potential (33%, 11 out of 33) or did not respond at all (24%, 8 out of 33). This distribution did not significantly change in 3-month-old organoids (33 neurons recorded, 11 with repetitive action potential, 11 with single action potential and 11 not responding). However, in 4-month-old organoids, most of the cells generated repetitive action potentials in response to depolarizing current injection (83%, 15 out of 18 cells). At the same time, rheobase did not significantly change with age of the neurons (2- to 4-months old organoids: 33.2±4.0 pA; 33.6±4.0 pA; 36.2±5.4 pA). These data indicate that neurons fully develop the ability to generate repetitive action potentials and switch from neonatal to more mature-type of electrical activity only by the age of four months that is consistent with human developmental timeline (24 gestational weeks; Silbereis et al., 2016).

Spontaneous neuronal activity is an important step in neuronal development. In immature cortex and hindbrain, the first spontaneous electrical activity is in the form of plateau-like depolarization, driven primarily by intracellular calcium fluctuations (Garaschuk et al., 2000; Gust et al., 2003). In the whole-cell patch clamp experiments, these oscillations can be seen as long-lasting fluctuations of the membrane potential (plateau-like voltage transients; Garaschuk et al., 2000; Allene et al., 2008; Dehorter et al., 2011). At the later stages, this plateau-like activity is substituted by faster action potentials. In the developing organoids, we observed a similar phenomenon. In 2- and 3-month-old organoids, many neurons displayed plateau-like voltage transients (Fig. 4C, 16 of 37 cells in 2-month-old organoids and 15 of 33 in 3-month-old organoids). However, in older (4 months) organoids almost all of the active cells displayed spontaneous action potentials (7 of 8 cells; 20 in total), and not voltage transients (1 of 20 cells)(Fig. 4E).

A similar developmental pattern can be seen when neuronal activity is detected as fluctuations of intracellular Ca^2+^. We detected calcium oscillations as early as at 2 weeks post-aggregation. However, Ca^2+^ transients were rare and could not be inhibited by TTX (tetrodotoxin, voltage-gated sodium channel blocker) (Supplemental Fig. 3A, B), consistent with mouse early hindbrain development (Gust et al., 2003). Two weeks after forced differentiation (1.5-month-organoid) the number of cells exhibiting spontaneous calcium transients markedly increased and up to 66.6±14.6 % (mean ± SD, n=7, 3 independent batches; custom R script, Supplementary code) of activity could be inhibited with TTX treatment (Fig. 4G; Supplemental movie 2). In 2-month-old organoids, some cells exhibited plateau-like calcium oscillations (Fig. 4F), consistent with electrophysiological data (Fig. 4C), which disappeared with the further development. Calcium waves pattern was diverse but some of the fluorescent traces showed a regular repetitive pattern (Fig. 4F; Supplemental Figs 3D & 4) that might be indicative of pacemaker activity, consistent with the idea that hindbrain can be one of the places hosting central pattern generator cells driving synchronous waves of activity in developing brain (Luhman et al., 2016).

Most organoids presented synchronous calcium waves after 15 days of forced differentiation (Supplemental Fig. 3G, H; Supplemental movie 3), which is consistent with *in vivo* published studies on mouse brain development (Garaschuk et al., 2000; Gust et al., 2003; Thoby-Brisson & Greer, 2008). Using a custom R script we calculated that a percentage of all possible pairs combinations of measured cells demonstrating correlated waves was 9.1±5.8 (mean±SD, n=11, 3 independent batches; Supplementary code). To determine the nature of the synchronization we inhibited electrical transmission by a broad spectrum connexins and pannexins inhibitor carbenoxolone (CBX). It had no effect on calcium transients nor on their synchronization (Supplemental Fig. 3C, D). The lack of CBX effect on local network bursts (synchronous activity) while inhibiting long-range propagation episodes was described before in the developing spinal cord of rodents (Hanson & Landmesser, 2003). Due to the lack of CBX effect on calcium transients it is likely that synchronization of cellular activity was synaptically mediated (Fig. 2F, G; Fig. 5).

Installation of synaptic network is one of the highlights of maturing nervous tissue. We further evaluated whether development of synaptic connections goes in parallel with maturation of the active membrane properties of the neurons. Both excitatory and inhibitory post-synaptic currents were already detected in two-month-old organoids (Fig. 5A-F) that is also consistent with human developmental timeline, considering that organoids were derived from already pre-patterned neural stem cells. Approximately 36% of patched cells in two-month-old organoids (9 of 25 cells) exhibited excitatory currents, and this number gradually increased (up to 65% in four-month-old organoids, 11 out of 17 cells). Neither the frequency of glutamate currents nor their amplitude changed significantly over time. The inhibitory synaptic network had already been set at 2 months of age - we observed that that nearly 75% of the neurons were synaptically connected irrespective of the age of organoids. However, the frequency of inhibitory currents increased significantly in the interval from two to three months of age. We also indirectly assessed the formation of synaptic contacts by transducing hNESCs with a lentivirus expressing GFP under human *Synapsin* promoter control (Tseng et al., 2018). After 2 months of differentiation the neurons, derived from the labelled cells, already demonstrated a robust expression of GFP that allowed us to visualize extensive neuronal processes (Fig. 5G; Fig. 7C). Taken together, we demonstrate that the sequential developmental events, shown in brain organoids *in vitro*, resemble the process of activity-driven synaptic maturation, described for neonatal rodent brain. Our data suggest that developing human neurons, cultured in 3D system, may be thus used as *in vitro* model for studying human synaptic development.

**Figure 5.**
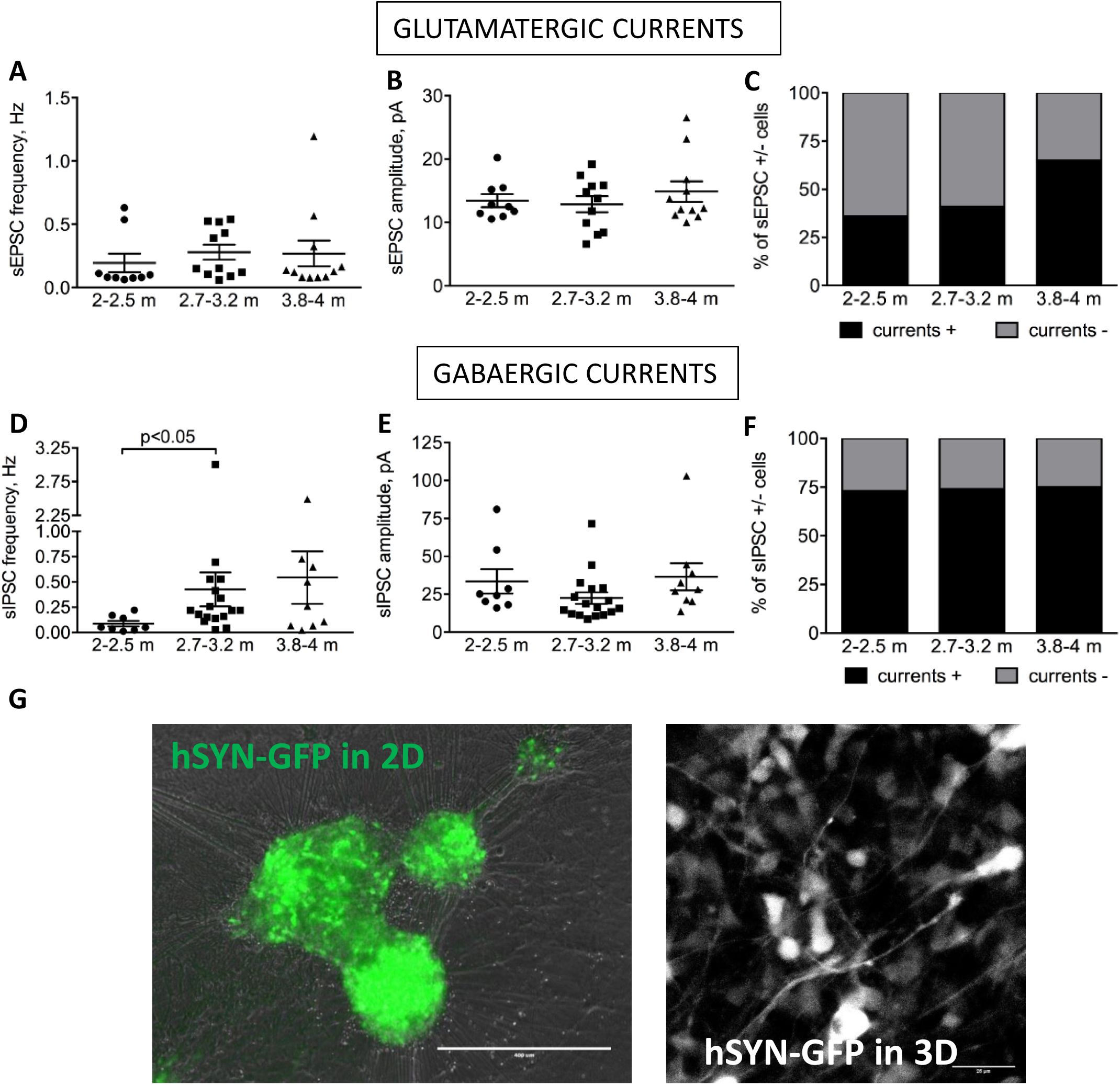
Excitatory and inhibitory synaptic currents in 2 to 4-month-old organoids (**A-F**). Frequency of spontaneous excitatory postsynaptic currents (sEPSC) (**A**), their amplitude (**B**) and the percentage of cells exhibiting them (**C**) do not change over time. Frequency of spontaneous inhibitory postsynaptic currents (sIPSC) (**D**) is significantly increased in 3-month-old organoids, while sIPSC amplitude (**E**) is not (*p<0,05, one-way ANOVA). **F.** Inhibitory synapses are present on nearly 75% of neurons and do not change significantly over time. **E**. Expression of human SYNAPSIN-GFP reporter in hNESC-derived neurons (differentiated for 17 days on Matrigel in two dimentions – 2D) and in 2-month-old organoids. Months – (m).

**Figure 6.**
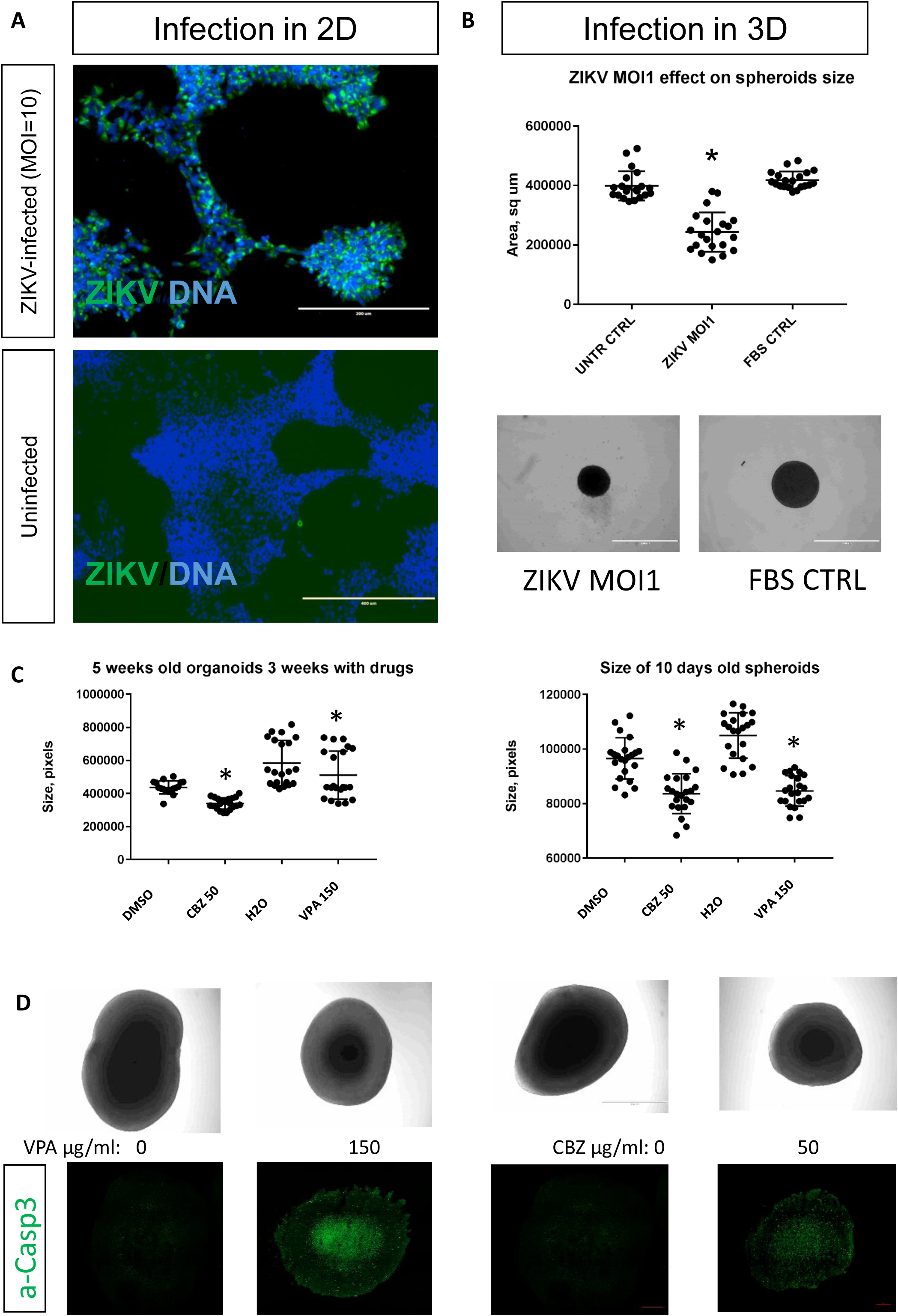
Effects of Zika virus infection and anti-convulsant drugs on organoids growth. **A**. Infection of hNESC in two dimensions (D) for 7 days. **B**. Infection of two-days-old hNESC spheroids in 3D. **C**, **D**. Effect of anti-epileptic drugs carbamazepine (CBZ) and valproic acid (VPA) on young spheroids growth (left) on older ones (right and also D, upper panel). **D**. CBZ and VPA increased cleaved caspase-3 staining in treated organoids. *p<0,05, Mann-Whitney U test.

**Figure 7.**
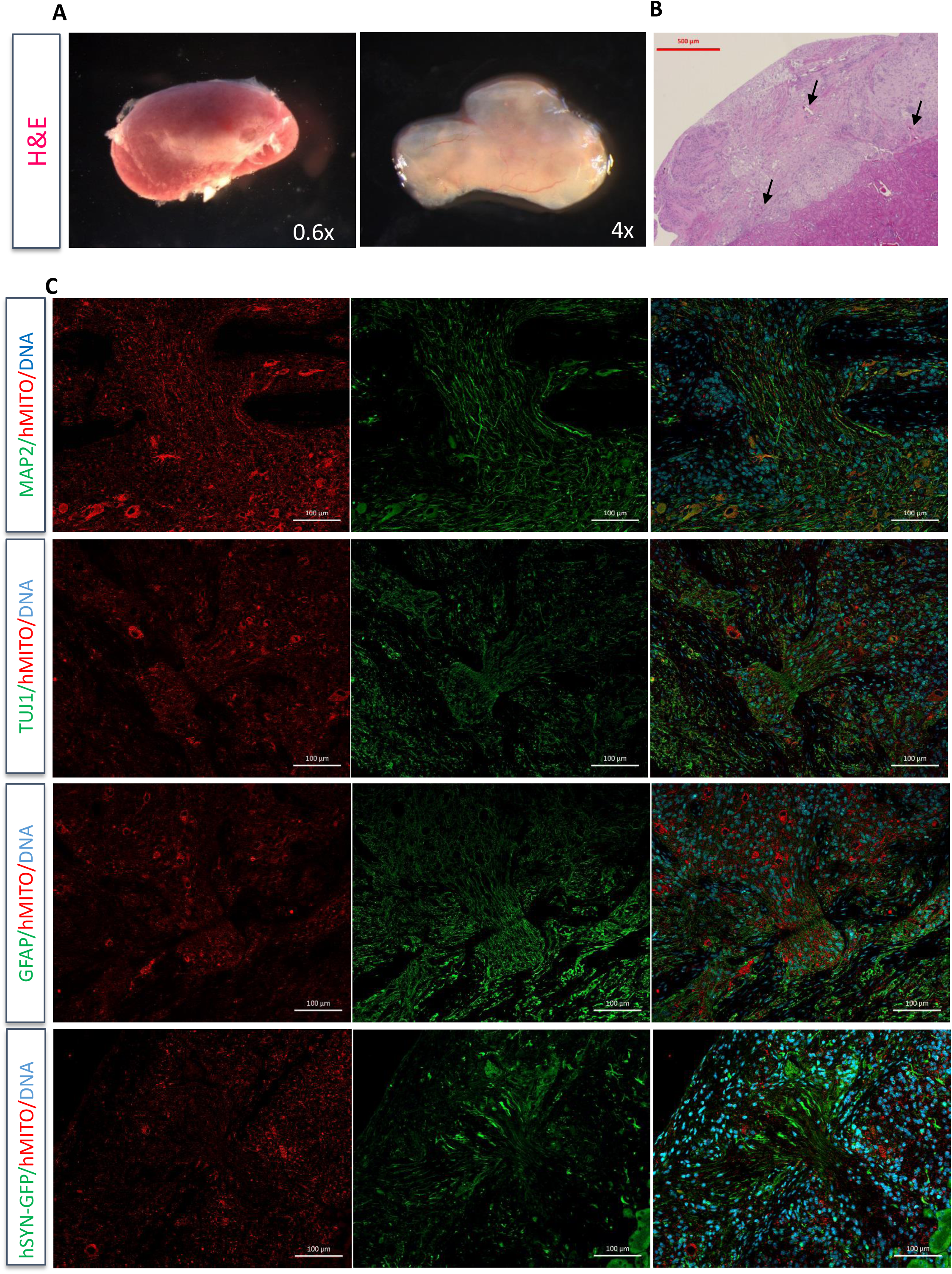
Transplantation of hNESC spheroids under the kidney capsule. **A**. Vascularisation of a graft. **B**. Hematoxylin and eosin staining of the graft. Blood vessels are marked with black arrows. **C**. Graft staining with neuronal (tubulin β-III - TUJ1; human SYNAPSIN-GFP reporter – hSYN-GFP) and astrocytic markers combined with staining of human antigen (hMITO). DAPI – stains DNA.

### Applicability of hNESC and brain organoids for disease modeling

To evaluate the applicability of caudalized hNESC and brain organoids for disease modeling we first tested whether Zika virus (ZIKV) can infect hNESC and hNESC-derived organoids. ZIKV is a mosquito-borne RNA-virus in the family *Flaviviridae,* causing microcephaly, among other central nervous system abnormalities, in children born to mothers infected in the first trimester of pregnancy. We found that hNESC can be effectively infected by ZIKV (Fig. 6A), as evidenced by immunofluorescence staining in 2D hNESC cultures. We have also observed that very young orgnoids (infected 2 days after the aggregation) produce viral particles (as measured by the increase in viral genome copy number in the supernatant; data not shown) and could efficiently be killed by ZIKV (MOI = 1) within a few days (Fig. 6B) that is consistent with published findings on forebrain neural stem cells (Qian et al., 2016). Importantly, this is a first demonstration, to the best of our knowledge, that caudalized hNESC could be effectively killed by ZIKV.

### Anticonvulsants reduce growth of brain organoids

To evaluate the applicability of caudalized brain organoids for drug testing we evaluated the effects of anticonvulsants of brain organoids growth. Pregnancy related epilepsy can affect up to 1% of women. Earlier epidemiological studies have linked anti-consultants like valproic acid (VPA) and carbamazepine (CBZ) to the increased risk of microcephaly in children born by mothers with epilepsy (Holmes et al., 2001; Jentink et al., 2010). We found that VPA and CBZ at concentrations 150 and 50 µg/ml, respectively, affected growth of brain organoids (Fig. 6C) by inducing apoptosis in neural progenitors as revealed by caspase-3 staining (Fig. 6D). The growth of both very young organoids (2 days after aggregation) and more mature ones (2 weeks) was significantly affected by VPA and CBZ compared to water and DMSO controls, respectively. The younger spheroids were treated for one week with anti-convulsants whereas older ones were treated for 3 weeks. This finding provides a mechanistic explanation to the teratogenic effects of the drugs observed in epidemiological studies.

## *In vivo* transplantation of brain organoids

Long-term maturation of hNESCs derived neurons requires a better availability to nutrients and oxygen due to organoids size and the lack of vasculature *in vitro*. In order to further improve maturation of neurons for functional studies we tested whether hNESC would survive and differentiate *in vivo.* We transplanted freshly aggregated hNESC (3-6 days post aggregation; 2-4 million hNESC in total per animal) under the kidney capsule of immunocompromised mice, the location that is easy to monitor and the one that provides adequate vascularization for transplanted cells (Saarimäki-Vire et al., 2017). In some experiments, we used hNESC transduced with lentiviral vectors expressing GFP under control of human *Synapsin* promoter. Two months after transplantation we found that the grafts survived and expanded, likely due to the support provided by the vascularization (Fig. 7A, B). The neurons in the grafts were MAP2- and GFP-positive and formed an extensive neuronal network that is intermingled by S100β-positive astrocytes (Fig. 7C). We found that some neurons remained spontaneously active even after 5 months post transplantation (Supplemental movie 4). This opens an avenue for modeling diseases and drugs effects on patient-specific or genome-engineered cells in humanized mouse model in the context of the entire organism and systemic drug administration.

## Discussion

The discovery of cell reprogramming and brain organoids opened up a fascinating opportunity to study living human neurons on molecular, single cell and local network level (Takahashi et al., 2007; Eiraku et al.,2008; Lancaster et al., 2013). However, due to a great diversity of neuronal cell types, it is important to clearly determine what region of the brain, type of the local network and neuronal cell types were modelled in a given protocol. Several brain regions, including cortex, hippocampus, striatum and *substantia nigra*, were already modelled using different approaches (Paşca, 2018). Here, we aimed to model hindbrain development.

During embryonic development, regionalization of the future brain is directed by concentration gradients of the signal molecules and by localized expression of transcription factors. In our protocol, we initiated the development of hNESCs by dual-SMAD (TGFβ and BMP signaling) inhibition and simultaneous induction of WNT and Hedgehog pathways, which stimulated proliferation, caudalization and, partially, ventralization of these neural stem cells. Protocols that lack induction of WNT and Sonic Hedgehog result in FOXG1-positive prefrontal cortex neurons (Chambers et al., 2009). Accordingly, when we tested the expression pattern of the main transcription factors, we found a high expression of *PAX6, MSX1* and *HOX* genes and low or virtually no expression of *NKX2.1, FOXG1* and *PAX7*. Such pattern corresponds to early development of myelencephalon – the developing brain area, which forms *medulla oblongata* in the adult brain.

Further differentiation of hNESCs into neurons in organoids was accomplished according to the published protocols. However, due to initial regionalization of neuronal progenitors, types of the neurons and their functional and morphological properties largely resemble rodent RVM. RVM and *medulla oblongata* are not only a relay station for pyramidal tract, located in medullar pyramids, this region also contains nuclei of cranial nerves with cholinergic neurons, and reticular formation, containing serotonergic, GABAergic and glutamatergic neurons (Morrison et al., 1991, Winkler et al., 2006; Stornetta et al., 2013). Using several approaches, we detected all these types of neurons in brain organoids produced with our protocol. Neuronal electrical properties and the shape of the dendritic tree resembled those of rodent serotonergic neurons in RVM (Table 1; Scott et al., 2005; Winkler et al., 2006; Lu et al., 2016). Moreover, in the mature organoids around 30 % of neurons fire spontaneous action potentials with the frequency of 2 Hz – similar to serotonergic neurons (Lu et al., 2016). These results further indicate the hindbrain regional identity of brain organoids.

Human developmental timeline is drastically different from that of a mouse, but most structural and functional hallmarks are present, nevertheless. In both species the synchronous spontaneous activity of neurons in cranial and spinal nerves (afferents of hindbrain and spinal cord) that drive embryonic motility appear early in development. At mouse embryonic day (E) 9.5 the motoneurons in hindbrain first exhibit TTX-insensitive, low-frequency, long-duration uncoordinated spontaneous activity that becomes synchronized and TTX-sensitive at E11.5 (Gust et al., 2003). Likewise, neurons of pre-Bötzinger complex, controlling respiratory rhythm, are synchronized at E15 (Thoby-Brisson & Greer, 2008). Serotonergic neurons are located in *raphe nuclei* and from medulla they initiate descending projections into emerging cervical spinal cord by E12.5 (Fenstermaker et al., 2010). In humans, the fetal breathing movements (FBMs) are first detectable at 10-12 weeks of gestation. At the start of third trimester (28 weeks) FBMs become more regular (Dawes et al., 1984; Greer, 2012), suggesting synchronization of neuronal activity. Roughly at the same time neonatal type of the network activity starts to appear in cortical areas after first thalamo-cortical pathways were established (>24 weeks; Iyer et al., 2015).

One-month-old organoids still contain neuronal progenitors, and postmitotic neurons start to massively appear after forced differentiation induction. This coincides with expression of synapsin, appearance of the first dendrites and excitatory and inhibitory synaptic currents. The neurite outgrowth and initial synapse formation do not require electrical activity of the cell in the form of action potentials (Cohen-Cory 2002, Varoqueaux et al., 2002, Verhage et al., 2000). Indeed, in two-month-old organoids only few cells were able to fire repetitive action potentials, but around half of the cells display synaptic currents, meaning that they are already interconnected into a functional synaptic network. The full maturation of the action potential firing happened only after four months after aggregation. Also, four-month-old neurons fired action potentials spontaneously, which correspond to the ability of hindbrain neurons to regulate the fetal breathing at six months of development of the human fetus (Greer, 2012).

Although synapses can be formed without action potentials, neuronal activity is essential for further maturation of neuronal circuits. During early development, the network neuronal activity evolves from long-lasting calcium oscillations and action potential-based network bursts to the adult-type action potential firing (Kerschensteiner, 2014; Luhmann et al., 2016). Different types of neonatal neuronal activity have been shown in various brain regions, including retina, hippocampus and cerebral cortex (reviewed e.g. in Blankenship and Feller, 2010; Moody and Bosma, 2005). Similar types of the neonatal neuronal activity have been recently demonstrated for organoids of the forebrain identity (Trujillo et al., 2019). Neurons in developing hindbrain also pass this developmental step, and the neonatal activity seem to be important in maintaining respiratory rhythm (Chatonnet et al., 2002). Here, in 2-3-month old organoids, we were able to record early-type spontaneous activity both on the level of intracellular calcium oscillations and as prolonged variations of membrane potential, or voltage transients. This activity pattern resembled slow calcium transients, some synchoronized, characteristic of developing rodent brain (Garaschuk et al., 2000; Gust et al., 2003; Allene et al., 2008; Dehorter et al., 2011). These long-term oscillations are substituted by regular action potential firing in older organoids, likely reflecting the maturation of the neuronal network.

Taken together, our data show that hindbrain organoids model the early steps of neuronal development, and four-month-old preparations likely correspond to the developing hindbrain in the third trimester of gestation period. Therefore, these organoids may serve as a promising model of the normally developing human brain and of that in a pathological condition. To test the latter, we treated growing organoids with Zika virus and two types of anticonvulsants, and, in accordance with the published clinical features, in all cases observed decrease in the size of organoids and neuronal death. Zika virus targets neuronal progenitors, causing severe microcephaly, and this effect was already modelled in human brain organoids (Gabriel et al., 2017; Watanabe et al., 2017) but not of the caudal origin. Antiepileptic drugs target the neuronal processes, important both for propagation of seizures and normal maturation of the synaptic connections, thus severely affecting the early brain development (Ikonomidou and Turski, 2010). For example, prenatal administration of valproic acid is the established model of autism spectrum disorder in rodents (Nicolini and Fahnerstock, 2018). To our knowledge, this is the first study showing the teratogenic effects of these anticonvulsants in human brain development model. Due to an ease to produce highly uniformed hNESC-based spheroids in 96-well format and due to a straightforward readout (diameter) this system could be further utilized for drug screening.

The presented model of hindbrain organoids could hopefully serve for other disease-related studies, such as modelling brainstem glioma invasion in this relevant human tissue (Bian et al., 2018), testing drugs in an individualized manner (Pemovska et al., 2013), or modelling neurodevelopmental disorders, e.g. neonatal seizures (Pospelov et al., 2016), and of mitochondrial diseases with atrophic medulla (Lonnqvist et al., 1998).

## METHODS

### Human samples and the use of animals

Human iPSC lines used in this study were generated after informed consent approved by the Coordinating Ethics Committee of the Helsinki and Uusimaa Hospital District (no. 423/13/03/00/08). H9 human embryonic stem cell line was purchased from WiCell. Animal care and experiments were approved by the National Animal Experiment Board in Finland (ESAVI/9978/04.10.07/2014). NSG (Jackson Laboratories; 005557) male mice, aged 3–12 months, were used for this study.

### Derivation of hNESCs

Undifferentiated human pluripotent stem cells (H9 ESCs line, HEL24.3, HEL47.2 human iPSCs lines; Balboa et al., 2017; Weltner et al., 2018)) were cultured on Matrigel (BD Biosciences)-coated plates in E8 medium (Life Technologies) and passaged with 5 mM EDTA (Life Technologies). One day before differentiation induction cells were plated in a density 1x10^6^ per 35 mm Matrigel-coated plate. On the next day (D0; Fig. 1A) cells were treated with 10 µM ROCK inhibitor (Selleckchem) for 5 hours in fresh E8 media and dissociated with StemPro Accutase (Thermo Fisher) to a single cell suspension by 5 min incubation at 37°C.

After dissociation cells were transferred to ultra-low attachment 6-well plates (Corning) and cultured in 2 ml of E6 media (Thermo Fisher), containing 10 µM SB431542, 100 nM LDN-193189, 3 µM CHIR99021 and 0.5 µM Purmorphamine (Selleckchem). The ROCK inhibitor Y-27632 in concentration 5 µM (Selleckchem) was present in the media for the first 48 hours after dissociation.

On day 2 the formation of the expanded neuroepithelium was assessed, and the media was changed to N2B27 medium (1:1) DMEM-F12/Neurobasal media; 1% NEAA, 0.5% GlutaMAX, 1% N2 Supplement, 2% B-27(without vitamin A) supplement (Life Technologies), 200 µg/ml Heparin, 2 mM L-Ascorbic acid (Sigma-Aldrich), containing 10 µM SB431542, 100 nM LDN-193189, 3 µM CHIR99021 and 0,5 µM Purmorphamine. On day 4 after aggregation SB431542 and LDN-193189 were removed from the media.

On the day 6 cell the aggregates were dissociated with StemPro Accutase (Life Technologies) into single cell suspension and plated on Matrigel (BD Biosciences)-coated plates. After 24 hours of plating 2.5 ng/ml bFGF (PeproThech) was added to the media. The final culture media for hNESCs expansion was composed of N2B27 media supplemented with 3 µM CHIR99021, 0.5 µM purmorphamine, and 2.5 ng/ml bFGF. Cells were routinely cultured on Matrigel-coated plates and split with Accutase after reaching a confluency at 1/10-1/20 ratio. The media was changed every other day.

### Derivation of brain organoids from hNESCs

For the generation of brain organoids hNESCs were dissociated to a single-cell suspension and aggregated in 100 µl of the standard culture media in round-shaped ultra-low attachment 96 well-plates (Corning) in density 2.5x10^3^-1x10^4^ cells per well. In some experiments we allowed spontaneous aggregation of NESC in rotating (95 rpm) 35 mm plates. The ROCK inhibitor Y-27632 in concentration 5 µM (Selleckchem) was present in the media for the first 48 hours after dissociation.

On day 6 after aggregation organoids were transferred on orbital shaker (95 rpm) and the culture media was changed to E6NB media that contains 1:1 E6/Neurobasal media (both from Thermo Fisher), 2% B27-supplement (containing vitamin A), 1% NEAA, 0.5% GlutaMAX (Thermo Fisher), 2% Human Insulin, 200 µg/ml Heparin, 100 µM L-Ascorbic acid (Sigma Aldrich) and Primocin (1:500; Invivogen).

On day 30 after aggregation the media was changed to the “forced differentiation media”, containing E6NB media supplemented with 0.4 mM dbcAMP (Sigma), 0.01 mM DAPT and 0,05 mg/ml human BDNF (PeproThech). In some experiments terminal differentiation media was added already 2 days post aggregation to decrease organoids size.

On day 60 the media was changed to E6NB media supplemented 0,01 mg/ml human BDNF (PeproThech), in which organoids were cultured until day 90 – day 120 depending on the purpose. The media was changed every other day.

The size of organoids was analyzed from images made with phase-contrast optical microscopy, using ImageJ software, the total area (pxl) was used as a computational parameter.

### Quantification of monoamines by HPLC with electrochemical detection

300 µl of the organoid media samples were filtered through Vivaspin filter concentrators (10,000MWCO PES; Sartorius, Stonehouse, UK) at 8600 g at 4 °C for 35 min. One hundred microliters of the filtrate was injected into chromatographic system with a Shimadzu SIL-20AC autoinjector (Shimadzu, Kyoto, Japan). The analytes were separated on a Phenomenex Kinetex 2.6 μm, 4.6 × 100 mm C-18 column (Phenomenex, Torrance, CA, USA). The column was maintained at 45 °C with a column heater (Croco-Cil, Bordeaux, France). The mobile phase consisted of 0.1 M NaH2PO4 buffer, 120 mg/L of octane sulfonic acid, methanol (5%), and 450 mg/L EDTA, the pH of mobile phase was set to 3 using H3PO4. The pump (ESA Model 582 Solvent Delivery Module; ESA, Chelmsford, MA, USA) was equipped with two pulse dampers (SSI LP-21, Scientific Systems, State College, PA, USA) and provided a flow rate of 1 ml/min. DA was detected using ESA CoulArray Electrode Array Detector with 12 channels (ESA, Chelmsford, MA, USA). The applied potentials for the channels were 1: 40; 2: 80; 3: 120; 4; 160; 5: 200; 6: 240; 7: 280; 8: 320; 9: 360; 10: 400; 11: 440 and, 12: 480 mV. DA was detected in the channel 2. The chromatograms were processed and concentrations of monoamines calculated using CoulArray for Windows software (ESA, Chelmsford, MA, USA).

### Metabolomic profiling

Untargeted metabolic profiling of hiPSC, hNESC and neural progenitors differentiated from hNESC (NPC) was performed using LC-MS with quadrupole time-of-flight mass spectrometry. The number of statistically significant peaks (p < 0.01; with fold change > 2) was 83 and 351 for pairwise comparison of hiPSC/hNESC and hiPSC/NPC, respectively (Supplementary table 2).

Cell samples (6 replicates of human induced pluripotent cells (hiPSC), neural stem cells (NSC), neural progenitor cells (NPC) and blanks) were lysed by adding 50 µl of MilliQ-water (Millipore, Molsheim, France) followed by 15 minutes sonication. 20 µl of cell lysate was transferred to a new deactivated insert and 10 µl of internal standard solution (PC(16:0 d31/18:1), PG(16:0 d31/18:1), LPC(17:0), 27-hydroxycholesterol-d6, verapamil, propranolol, palmitidic acid-d31, ibuprofen and pregnenolone-d4) and 100 µl of methanol was added to precipitate the proteins. Mixture was vortexed and incubated in ice for 30 minutes, followed by centrifugation (10 min, 10 000 rpm, room temperature). Supernatant was transferred to a new vial with a new deactivated insert. The total protein content of the homogenate was determined with NanoDrop 1000 (Thermo Fisher Scientific, Waltham, MA, USA) for normalization purposes.

LC-MS analysis was performed with Synapt-G2 quadrupole time-of-flight mass spectrometry (Waters, Milford, MA, USA) combined with Acquity UPLC (Waters, Milford, MA, USA). The instrument was controlled with MassLynx^™^ 4.1 software and data were collected in centroid mode using lock mass correction of leucine-enkephalin. Mass range was 100-1000 m/z, with scan time of 150 ms, applying extended dynamic range and sensitivity mode (resolution 10 000 FWHM). The MS operation parameters were: capillary voltage 2.5 kV, sampling cone voltage 30 V, extraction cone voltage 3 V, source temperature 150 °C, desolvation temperature 450 °C, desolvation gas flow 800 L/h and cone gas flow 20 L/h.

The UPLC reversed phase column was BEH C18 (2.1 mm x 100 mm, 1.7 µm) from Waters (Milford, MA, USA). Eluent A was 0.1% formic acid in MilliQ-water and eluent B 0.1 % formic acid in methanol. Linear gradient started from 5% of B and proceeded to 100% B in 15 minutes, kept there for 10 minutes followed by the re-equilibration step of 5 minutes. Total runtime was 30 minutes with the flow rate of 0.3 ml/min. Autosampler was cooled to 10 °C and column thermostated for 25 °C degrees. Injection volume was 5 µl with a partial loop injection mode.

Waters .raw data files were converted with DataBridge to netCDF files prior the data preprocessing, which was performed with MzMine 2 1. All features that were found from blank and did not show difference between one of the cell groups and blank (t-test, q>0.05) were deleted. Furthermore, metabolic features were normalized with internal standard and total protein content. Data analyses were performed with R version x64 3.0.2. utilizing metadar-package2. Zero values in the data were imputed with a half of the minimum value of the corresponding feature across all of the samples. In addition, data was scaled prior to the unsupervised principle component analysis with log2 transformation. ANOVA followed by Tukey’s HSD post-hoc test and FDR correction (Benjamini-Hochberg) was applied to compare metabolite differences between cell groups (Pluskal et al., 2011; Peddinti 2012).

### Proteomics

#### Sample preparation

Water was purified with Milli-Q water purification system (Millipore, Molsheim, France). Six biological replicates of human induced pluripotent cells (hiPSC), neural stem cells (NSC), and neural progenitor cells (NPC) were thawed, and the total protein content of the samples was measured with NanoDrop 2000 in triplicate measurements (Thermo Scientific, Waltham, MA USA). The cells were lysed by 15 min sonication in 8 mol/L urea (Amresco, Solon, OH, USA). Insoluble cell debris was removed by two 15 min centrifugations at 20000 G, after which the supernatant was diluted with water to < 1.5 mol/L urea. Disulphide bonds were reduced with 5 mmol/L dithiothreitol and the cysteine residues carbamidomethylated with 15 mmol/L iodoacetamide (final concentrations; Sigma-Aldrich, Steinheim, Germany), followed by adjusting the pH to ∼8 with 1 mol/L ammonium bicarbonate (Fluka, Buchs, Switzerland) and overnight digestion with sequencing grade modified trypsin (Promega, Madison, WI, USA).

The digested samples’ pH was adjusted to ∼2 with 10 % trifluoroacetic acid (Sigma-Aldrich) and LC-MS grade acetonitrile (Merck, Darmstadt, Germany) was added to the final concentration of 1 %. The samples were purified with C18 MicroSpin columns (The Nest Group, Southborough, MA, USA), followed by evaporation to dryness in a vacuum centrifuge and storage in -20 °C. Before the LC-MS analysis, the samples were resolubilized by 15 min sonication in 0.1 % trifluoroacetic acid, 1 % acetonitrile in LC-MS grade water.

#### LC-MS analysis

Approximately 6 μg of total digested protein was used for the LC-MS analysis, which was carried out as described in Loukovaara et al. 2015. Briefly, the tryptic peptides were separated with a 150 min gradient run on a ThermoEASY nLC II using C18 pre-column and analytical column. The detection was done with Thermo Orbitrap Elite MS in top20 data-dependent acquisition mode.

#### Data analysis

Protein identification and quantification were done with Andromeda and MaxQuant, respectively (Cox & Mann 2008; Cox et al. 2011, 2014). An up-to-date *Homo sapiens* UniProtKB/Swiss-Prot proteome was used as the library (The UniProt Consortium 2017). The mass tolerance was 4.5 ppm for MS1 and 0.5 Da for MS2, and the maximum of two missed cleavages were allowed. Identifications were filtered for 1 % FDR on both peptide and protein level. Before further statistical analysis, contaminant and decoy matches were filtered out. One sample (NPC_04) was removed as an outlier due to unsuccessful LC-MS run. Protein LFQ intensities were used for abundance comparison, and for statistical testing, zeros were imputed with noise normally distributed around the non-zero minimum LFQ intensity of each protein divided by 10. For statistics, ANOVA followed by Tukey’s HSD post-hoc test and FDR correction were applied, and proteins with FDR < 0.01 were considered significantly differentially expressed.

### Immunofluorescence

The adhesive cell cultures were fixed in 4% PFA for 15 min, permeabilized with 0,1-0.5% triton-X100 in PBS, blocked with UltraV block (Thermo Fisher) for 10 min and incubated with primary antibodies diluted in 0,1%Tween in PBS at 4 °C overnight. The plates were washed with PBS, incubated with secondary antibodies (AlexaFluor; Thermo Fisher) diluted in 0.1% Tween in PBS for 30 min.

Cerebral organoids were fixed in 4% PFA for 20 min, washed in PBS and incubated in 20% Sucrose in PBS at 4 °C overnight. Organoids were incubated in Tissue Freezing Medium for 20 min and subsequently frozen at -20 °C; 20 µm sections were obtained using cryotome. Sections were washed with PBS and stained similarly to adhesive cell cultures.

The list of Primary antibodies and dilutions are listed in Table S3.

### Histology/Immunohistochemistry

The kidneys were cut as two halves and fixed with 4 % PFA in RT overnight. After this, samples were transferred to cassettes and processed for tissue transfer. 5 μM sections were cut with microtome. The slides were deparaffinised. Antigen retrieval was performed by boiling slides either in 1 mM EDTA or 0.1 M citrate buffer. Blocking and incubation with primary and secondary antibodies were done as described for fixed cells above.

### Electrophysiology

Whole-cell patch-clamp recordings were done on living slices, prepared from organoids of different ages after aggregation. One or two organoids were mounted into low gelling temperature agarose block, and 300 μm-thick slices were made using a vibratome (Leica 1200) in ice-cold cutting solution of the following composition: 117 mM choline chloride, 2.5 mM KCl, 0.5 mM CaCl2, 1.25 mM NaH2PO4, 7 mM MgSO4, 26 mM NaHCO3, 15 mM glucose, and 95 % O2/5 % CO2. Slices were left to recover for 1 hour at 34^0^C in recording ACSF with concentration of MgSO4 elevated to 3 mM. After a recovery period, slices were transferred to the submerged recording chamber on the visually guided patch-clamp setup. Recordings were done at 30^0^C in ACSF of the following composition (mM): 124 mM NaCl, 3 mM KCl, 1.25 mM NaH2PO4, 1 mM MgSO4, 26 mM NaHCO3, 2 mM CaCl2, 15 mM glucose, and 95 % O2/5 % CO2. Whole-cell voltage clamp recordings were done with glass microelectrodes (4-6 MOhm), using Multiclamp 700A amplifier (Axon Instruments, USA), and digitized at 20 kHz. Data were collected with WinLTP 2.30 (WinLTP Ltd. and University of Bristol, UK) and analyzed in Clampfit 10.2 (Molecular Devices, USA) and Mini Analysis program 5.6.6 (Synaptosoft, USA). Meta-analysis was done in GraphPad Prism 6. Data are presented as per cents from the total number of cells per condition, or individual measurements and mean±SEM. Statistical significance of the results was estimated by Chi-square test or Mann-Whitney test.

*Passive and active membrane properties, as well as sEPSCs* were recorded with the filling solution, containing: 116 mM K gluconate, 15 mM KCl, 5 mM NaCl, 10 mM HEPES, 4 mM MgATP, 0.5 mM Na2GTP, 0.5 % biocytin, pH 7.2. *S*eries resistance, input resistance and capacitance were recorded in voltage clamp mode at -70 mV by injection of 5 mV voltage steps, and if the change in series resistance during the experiment exceeded 30 %, the recording was discarded. Resting membrane potential and spontaneous activity were recorded in current clamp mode without any background current injected. For recording of the evoked action potentials, baseline membrane potential was held at -70 mV, and depolarising current steps with the increasing increment of 10 pA were injected for one second. For recording of sEPSCs, cells were kept in voltage clamp mode at -70 mV. *sIPSCs* were recorded in voltage clamp mode at -70 mV in the presence of 20 μM NBQX and 5 µM L-689,560. For these recordings, cells were filled with solution of the following composition: 120 mM CsCl, 0.022 mM CaCl2, 4 mM MgCl2, 10 mM HEPES, 0.1 mM EGTA, 5 mM QX-314, 4 mM MgATP, 0.5 mM Na2GTP, 0.3 % biocytin, pH 7.2.

### Analysis of the dendritic tree

After electrophysiological recordings, slices were fixed in 4% PFA overnight, after which they were washed with 0.01M PBS and permeabilized with 0.3% Triton-X 100 in PBS (Sigma-Aldrich) for 2h. Alexa fluor 568 or 488 streptavidin conjugate (1:1000; Life Technologies) was added to the permeabilization solution and incubated for 2h. After staining, slices were mounted using Fluoromount mounting medium (Calbiochem).

Labelled neurons were imaged with LSM Zeiss 710 confocal microscope (EC Plan-Neofluar 20x/0.50 M27) with resolution of 1.69 pixels/μm and Z-stack interval of 2 μm. Images were traced using the Simple Neurite Tracer plugin (Longair et al., 2011), and files were processed by Sholl analysis (Sholl 1953) (Fiji Is Just ImageJ, NIH, Bethesda, MD). The center of analysis was defined by a point ROI at the cell soma, step size of radius 10 µm. Number of intersections across the distances were used for statistical analysis.

### Construction of transfer plasmids and lentiviral vectors production

A 448 nt fragment of human *SYN1* promoter was PCR amplified using Phusion high fidelity DNA polymerase (Thermo Scientific) and the following primers: BcuI_hSYN_for 5’-ATACTAGTAGTGCAAGTGGGTTTTAGGACC and EcoRI_hSYN_rev 5’-TGGAATTCGACTGCGCTCTCAGG, digested with FastDigect-BcuI/-EcoRI (Thermo Scientific), and cloned into pCDH-CMV-MSC-T2A-EGFP vector (System Biosciences) digested with the same restriction enzymes to obtain pCDH-hSYN-EGFP. Lentiviral vectors (LVs) were produced as described (Tseng et al., 2018).

### Quantitative RT-PCR (qRT-PCR) analysis

For gene expression analysis NESC were grown to various passages and quantitative RT-PCR (qPCR) was performed in 3 lines in duplicates.

RNA was extracted with NucleoSpin RNA II kit (Machery-Nagel RNA, 740948) and RT-reaction was performed as described previously (Toivonen et al., 2013). qRT-PCR reactions were prepared with 5x HOT FIREPol® EvaGreen® qPCR Mix Plus (no ROX) in a final volume of 20 μl, containing 50 ng of RNA retrotranscribed to cDNA and 5 μl of a forward/reverse primer mix at 2 μM each. The reagents were pipeted with Corbett CAS-1200 liquid handing system. The qPCR was performed as duplicates using Corbett Rotor-Gene 6000 (Corbett Life Science) with a thermal cycle of 95°C for 15 min, followed by 45 cycles of 95°C, 25 s; 57°C, 25 s; 72°C, 25 s, followed by a melting step. Relative quantification of gene expression was performed following the ΔΔCt method using housekeeping genes *Cyclophillin G* and *Gapdh* as endogenous loading control and a RT-reaction without template as a negative control. Exogenous positive control was used as a calibrator. Data was presented as a fold-change using normalization with expression level of undifferentiated H9 human embryonic stem cells cells.

### Calcium imaging

For calcium imaging, a concentration of 5 µM cell permeant Fluo-4 AM (Life Technologies) in a conditioned E6NB medium with 10 ng/ml BDNF was added and incubated for 30 min at 37 °C on an orbital shaker, following by media change in order to eliminate the dye excess. Whole organoids were transferred to imaging plates and mounted into Matrigel (Corning). Fluorescent images were acquired using a live cell recording with Zeiss LSM 880 confocal microscope, in the conditioned culture media at 37 °C, 5% CO2.

The images were analyzed using ImageJ software to set ROIs, presenting individual cell bodies, and measuring mean intensity across every frame of recorded time-series. The background fluorescence was subtracted and cells activity were presenting using ΔF/F0 parameter.

### Implantation into testicles and under kidney capsule

NOD-SCID-gamma (NSG) mice were housed at Biomedicum Helsinki animal facility, on a 12h light/dark cycle and food ad libitum.

To test a potential teratogenicity of hNESC we first transplanted increasing amounts of dissociated hNESC (0.5, 1 and 2 million) cells into testis of immunodeficient non-obese diabetic (NOD)-severe combined immunodeficiency (SCID)-gamma (NSG) mice to test the potential for teratoma formation of hNESC in the same manner we routinely perform evaluation of iPSC differentiation potential (Weltner et al., 2017). Two months after transplantation we found no evidence of teratoma formation (data not shown).

Approximately 5 million cells were transplanted as aggregates under the kidney capsule Aggregates were loaded in a PE-50 tubing connected to a needle and centrifuged to form a tight pellet inside the tubing. Aggregates-loaded tubings were kept on ice until transplantation. Mice were anesthetized with isoflurane. Small incision of about 1 cm was cut open in mouse left flank skin and underlying peritoneum. Left kidney was carefully pulled out and a small nick was practiced with a needle on the apical side of the kidney to create a minimal open in the kidney capsule. A glass rod was used to open a pocket under the kidney capsule, the PE-50 tubing was introduced carefully and the aggregates were slowly released by screwing a Hamilton syringe coupled to the tubing. The kidney capsule opening was sealed with a cauterizer. Both peritoneum and skin were stitched with surgical silk. Carprofen (Rimadyl, 5 mg/kg, sc, Prizer, Helsinki, Finland) and Buprenorphine (Temgesic, 0,05-0,1 mg/kg, sc, RB pharmaceuticals Lmt, Berkshire, Great Britain) were used as analgesics during the operation and in the following day.

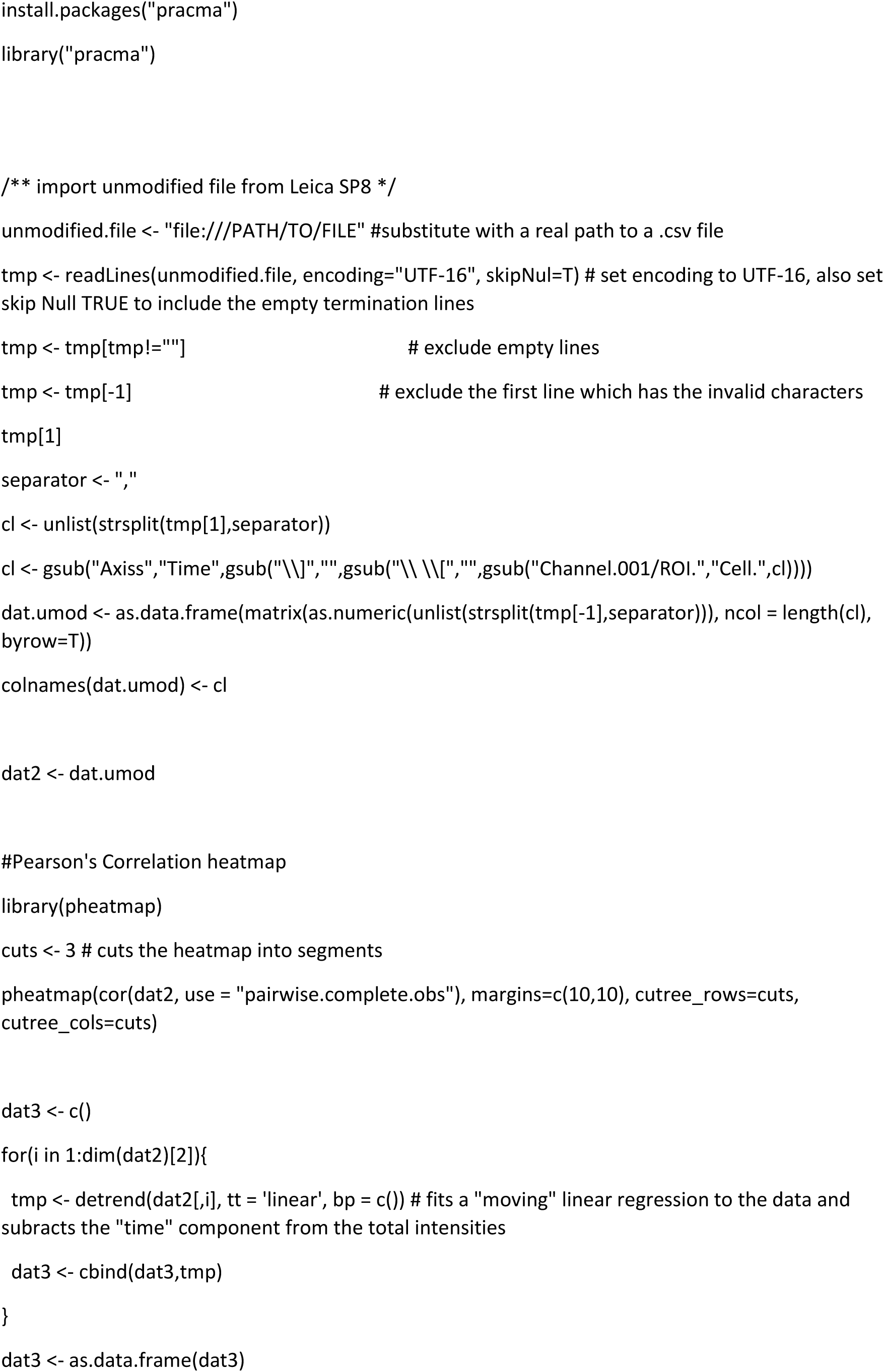
Custom R script.

## Acknowledgements

We are thankful to Heli Grym, Anni Laitinen, Agnes Viherä, M.Sc. Jarkko Ustinov, M.Sc. Anna Näätänen, and Eila Korhonen for an excellent technical assistance. We acknowledge Dr Ras Trokovic and Biomedicum Stem Cell Center for providing hiPSC lines. We thank Dr Evgeny Prjazhnikov and Prof. Claudio Rivera for help with establishing Ca^2+^-imaging assay; and Liliia Andriichuk for help with Ca^2+^ transients synchronization analysis. Dr Pavel Uvarov is acknowledged for critically reading the manuscript. We thank 3iRegeneration (Business Finland) and EDUFI for financial support. AD is supported by the Academy of Finland (grants #293392, #319195). RRG is supported by the Russian Science Foundation (grant N 19-75-30008). SK and OV thank Academy of Finland.

---

**Supplementary Figure 1. A.** Comparison of hNESC aggregation comparing previously published and updated protocol. Scale bar is 200 µm. **B**. Freshly plated hNESC colonies (passage 0) lack pluripotency marker OCT4 and endodermal marker GATA4 but almost uniformly express VIMENTIN (Vim). Some cells express MASH1/ASCL1 and doublecortin (DCX). **C**. Additional qRT-PCR charachterization of hNESC generated from HEL24.3 hiPSC line. **D**. Principal component analysis (PCA) and hierarchical clustering of proteomics data derived from hiPSC (HiPSC), differentiated for 3 days hNESC (NPC) and hNESC (NSC). **E**. PCA of untargeted metabomoics data. Labels are as in D.

**Supplementary Figure 2.** High performance liquid chromatography measurement of monoamines in conditioned media of organoids. 5HT – serotonin.

**Supplementary Figure 3. A.** Rare calcium transients in immature, 20 days post aggregation, spheroids that could not be inhibited by tetrodotoxin (TTX) (**B**). **C.** Maximum intensity Z-projection by SD (standard deviation) of a more mature, day 15 post differentiation induction with γ-secretase inhibitor DAPT, organoid. **D.** Transients were unaffected by carbenoxolone (CBX). Data on A, B are expressed as ΔF/F0 over 34 min interval. Data on D are isolated traces of selected cells over 5 min interval (extracted from a 31 min recording). **E-H**. Synchronization of Ca^2+^ transients. **E.** A still with regions of interest from recording with Fluo4 dye of a 2-month-old organoid (Supplemental movie 3) and corresponding calcium transients (**F**). Correlation analysis of these calcium transients presented as a heatmap (**G**) and as a correlation plot (**H**).

**Supplementary Figure 4.** Variability in calcium transients in a 2-3 months old organoids

**Supplementary Movie 1.** A 1-month-old organoid (left) and a 1.5-month-old organoid (2 weeks after forced differentiation) on the right. MAP2 in green and SOX2-tdTomato (Balboa et al., 2017) is red.

**Supplementary Movie 2.** Calcium imaging (Fluo4) in brain organoids 15 days post differentiation induction before (left) and after tetrodotoxin (TTX; right) treatment. The movies were recorded for 30 min and then sped up 40 times. Lookup table (ImageJ 16 colors) was applied to visualize changes in fluorescence intensity.

**Supplementary Movie 3.** Calcium imaging (Fluo4) in 3-month-old organoid. The movie was recorded for 3 min and then sped up 40 times. Lookup table (ImageJ 16 colors) was applied to visualize changes in fluorescence intensity.

**Supplementary Movie 4.** Calcium imaging (Fluo4) in an extracted 5-month-old graft. The movie was recorded for 3 min and then sped up 40 times. Lookup table (ImageJ 16 colors) was applied to visualize changes in fluorescence intensity.

