## Supplemental figures and Table 2 for "Maturation of Neuronal Activity in Caudalized Human Brain Organoids"

Supplemental Figure 1

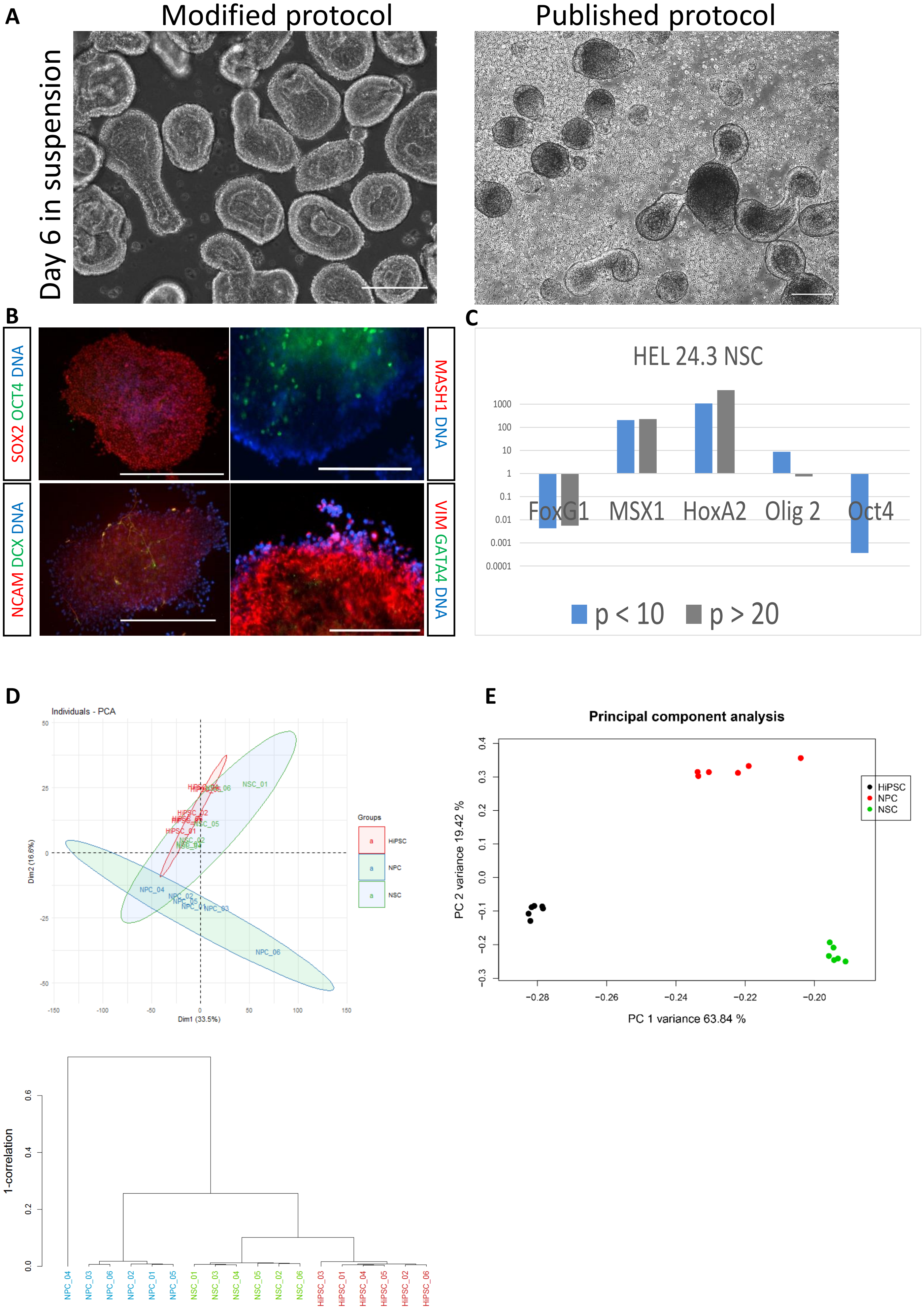

### Supplemental Figure 2

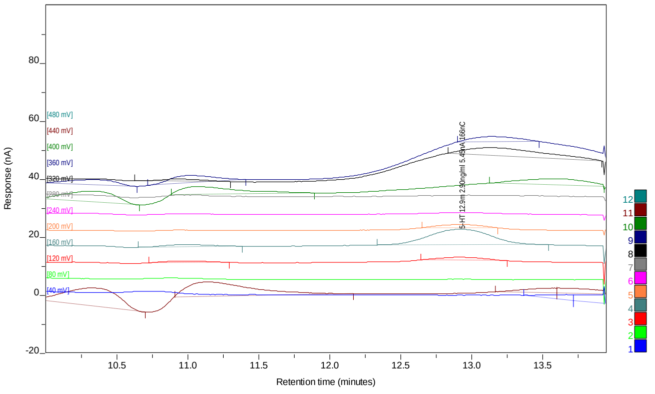

### Supplemental Figure 3

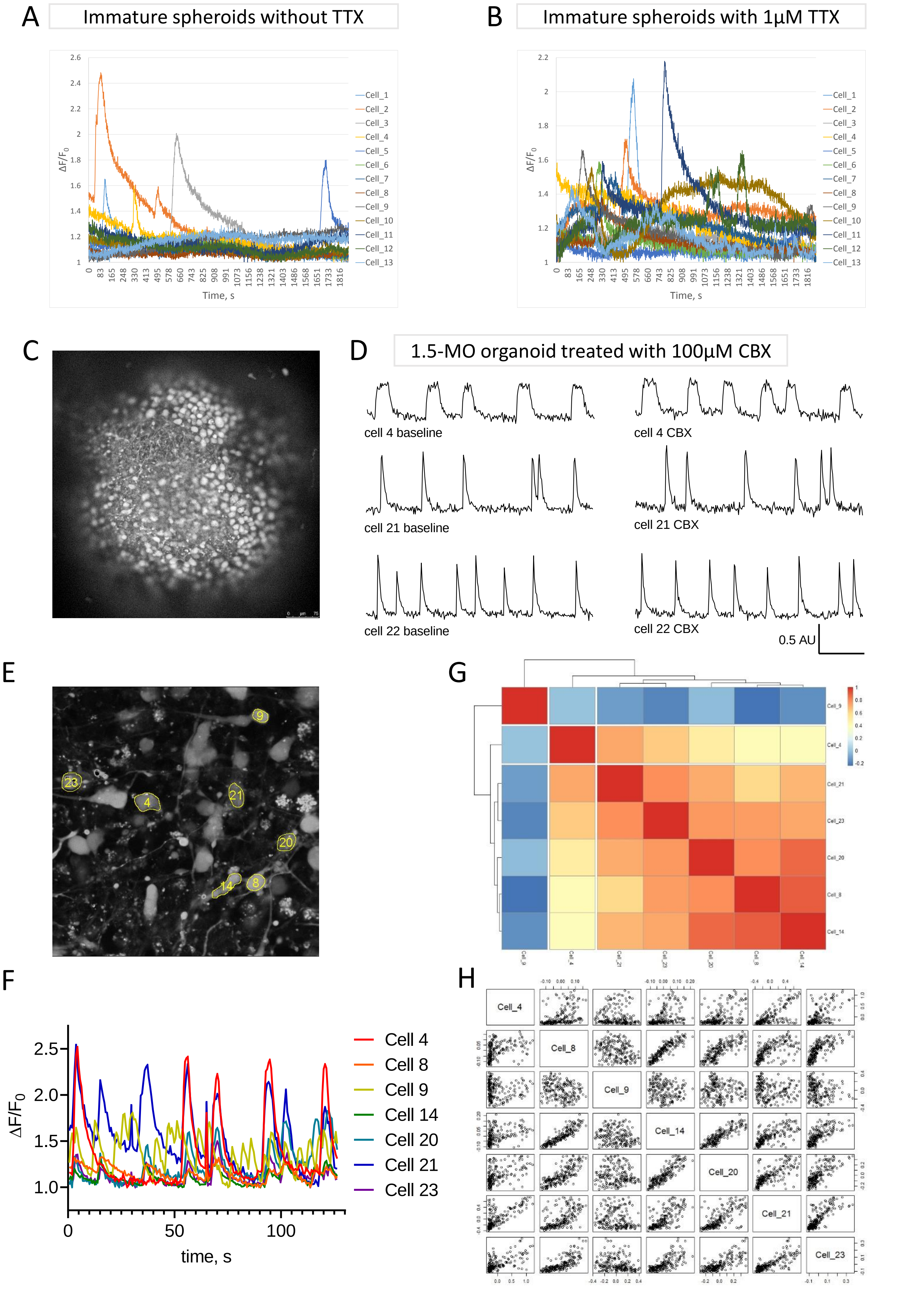

### Supplemental Figure 4

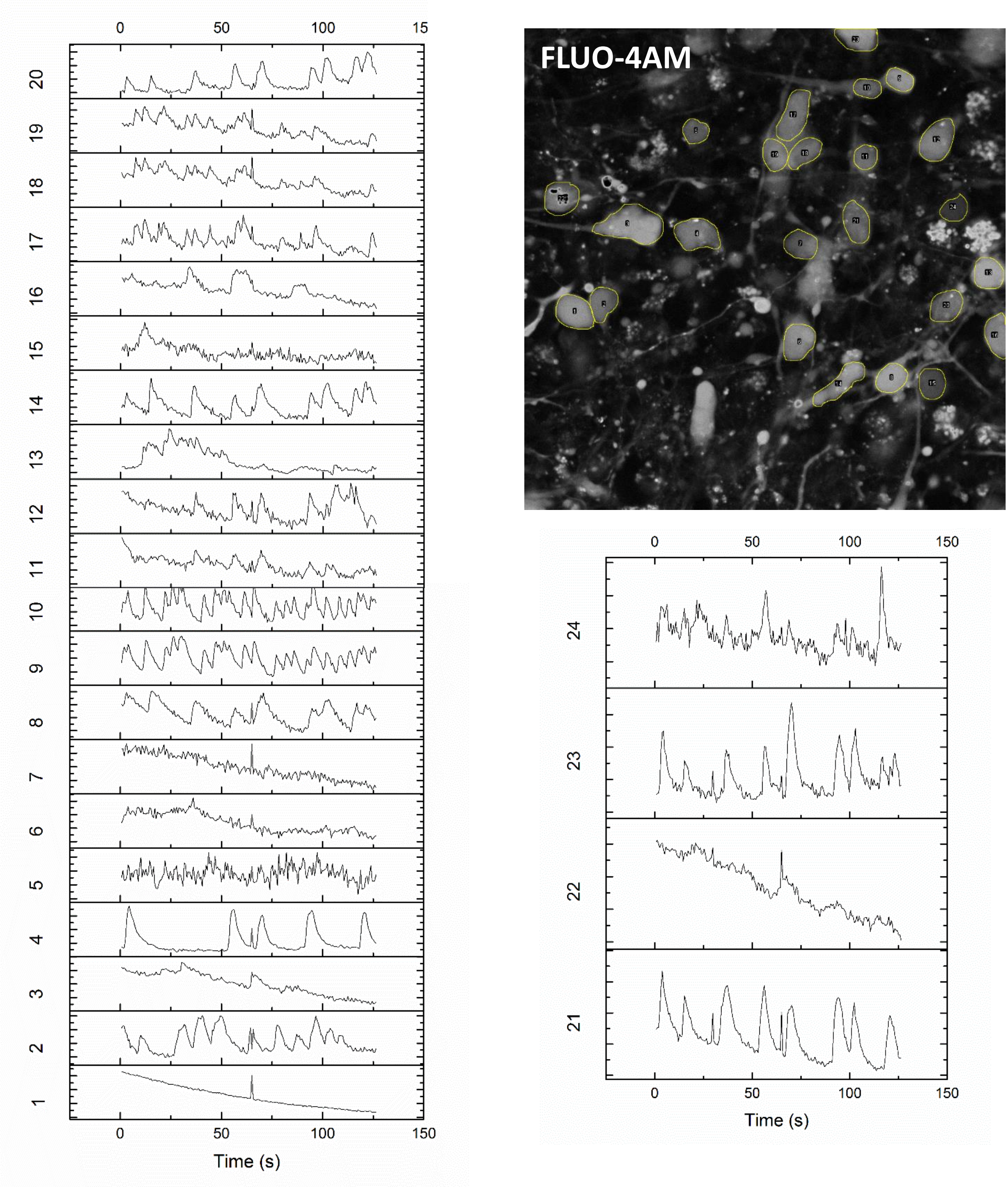

### Supplementary table 2

| Pairwise comparison of statistically important peaks between cell types | | | |
|---|---|---|---|
| Neural stem cells | NSC/hiPSC | NPC/hiPSC | NPC/NSC |
| Number of statistically significant peaks (q<0.01), with fold change > 2 | 106 | 376 | 128 |
| Number of statistically significant peaks (q<0.01), with fold change <0.5 | 304 | 44 | 34 |
| **Number of statistically significant peaks (q<0.01), with fold change > 2 or <0.5 in total** | **410** | **420** | **162** |
| *ANOVA followed by Tukey´s HSD post-hoc test with FDR correction (Benjamini-Hochberg) | | | |
